# Organoids serve as viable *in vitro* model for functional precision medicine for mesonephric-like adenocarcinoma of the ovary

**DOI:** 10.64898/2026.04.09.717457

**Authors:** David Holthaus, Huy Duc Le, Laura Matzner, Franziska Kellers, Christoph Rogmans, Vincent Winkler, Lorenz Bastian, Stephanie Fliedner, Jörg-Paul Weimer, Hauke Busch, Tim Mandelkow, Björn Konukiewitz, Nicolai Maass, Marion van Mackelenbergh, Ibrahim Alkatout, Dirk O. Bauerschlag, Nina Hedemann

## Abstract

**Background:** Mesonephric-like adenocarcinoma has been recently classified as a rare type of ovarian carcinoma. Description of these tumours have been rare and mostly covered in case reports. In some cases, molecular characterization by sequencing has been employed for guided therapy recommendations, however, functional chemosensitivity testing of targetable pathways using advanced *in vitro* cellular models such as organoids has not been reported so far. Here, we report on a case of ovarian cancer that was later identified as mesonephric-like adenocarcinoma at an advanced stage.

**Methods:** The tumour was characterized by molecular techniques including immunohistochemistry and whole-exome sequencing. At the same time, ovarian cancer organoids were established by adapting existing protocols for high-grade serous ovarian carcinoma. The organoids were subsequently used for functional *in vitro* chemosensitivity testing by treatment with standard-of-care chemotherapeutics cisplatin, paclitaxel, and the Poly (ADP-Ribose) Polymerase 1-inhibitor olaparib. Based on molecular characteristics, we also applied the inhibitor binimetinib, to target Mitogen-Activated Protein Kinase downstream of the KRAS Proto-Oncogene. Additionally, chemotoxicity testing with healthy fallopian tube organoids and high-grade ovarian cancer organoids was applied to determine the therapeutic window.

**Results:** Immunohistochemical analysis showed characteristic PAX8^+^, GATA3^+^, TFF1^+^, ER^-^, PR^-^, WT1^-^ staining while the sequencing revealed mutations in 31 genes of which KRAS G12V and DYNC1H1 G4072S were annotated as (likely) pathogenic. The tumour was mismatch-repair proficient. Tumour-derived organoids proved to be highly resistant to standard-of-care chemotherapeutics cisplatin, paclitaxel, and olaparib, but sensitive to inhibition by binimetinib, which aligned well with the molecular characteristics. Direct comparison to healthy fallopian tube organoids and high-grade ovarian cancer organoids confirmed low cytotoxic potential underlining a feasible therapeutic window for binimetinib.

**Conclusions:** For the first time, we show that existing protocols for high-grade serous ovarian carcinoma can be used for the generation of organoids derived from mesonephric-like adenocarcinoma. These organoids could be used as an essential tool for functional precision medicine purposes. This functional data could be applied as an additional layer for molecular tumour boards diagnostics by supporting molecular datasets and even identify targetable pathways beyond genetic variations, thus offering novel therapeutic options particularly for rare and aggressive tumours.

## Background

Recently, a new type of ovarian carcinoma has been classified, the mesonephric-like adenocarcinoma (MLA) that accounts for less than 1% of all cases (1). MLAs represent a rare type of malignant neoplasm affecting the uterine corpus and ovaries. It is hypothesized that mesonephric duct remnants persist along the lateral walls of the cervix, vagina, uterine corpus and adnexa during embryogenesis. These duct remnants are believed to degenerate, and sometimes trigger malignant transformation (2, 3). In recent years another hypothesis concerning the pathogenesis of mesonephric-like adenocarcinomas has emerged from a more thorough examination of its molecular characteristics: Mesonephric-like adenocarcinomas have similar characteristics to Müllerian-type carcinoma with secondary mesonephric differentiation (1, 4). Despite its rare occurrences and the fact that they have been mainly described in case reports, mesonephric-like adenocarcinomas have been included in the updated World Health Organization Classification of Female Genital Tract Tumours in 2020 (5). Pors, Segura (3) recently presented the largest clinicopathological characterization of 25 cases of MLA: The mean age of disease onset was 61 years. Patients presented with symptoms of pelvic pain, postmenopausal bleeding and abdominal bloating. Furthermore, 39% of patients were diagnosed at an advanced stage (FIGO ≥ 2), with 42% of cases exhibiting disease recurrence, predominantly in distant sites (3).

Currently, there are no guideline recommendations for the treatment of MLA. As with other ovarian carcinomas, cytoreductive surgery with maximum tumour resection is generally recommended. The surgical procedure should include a total hysterectomy, bilateral adnexectomy, omentectomy and pelvic lymphadenectomy. This is, due to the lack of feasible alternatives, usually followed by adjuvant chemotherapy with carboplatin and paclitaxel, as with high-grade serous ovarian cancer (HGSOC) (6). A challenge here is that MLA is associated with a poorer prognosis than other ovarian carcinoma subtypes as it often exhibits chemotherapeutic resistance and a higher recurrence rate. It is therefore crucial to carry out an extended characterization of this tumour entity in order to evaluate new therapeutic approaches.

However, functional testing of chemotherapy, including standard-of-care (SOC) platin- and taxane-based therapeutics, using *in vitro* propagated cells has been missing for this particular tumour entity. Here, we describe a rare case of mesonephric-like adenocarcinoma of the ovary of which we have developed organoids that were used for functional *in vitro* chemoresistance determinations including chemotoxicity testing using benign fallopian tube epithelium alongside of standardized molecular characterization by whole exome sequencing (WES) and immunohistochemistry (IHC) in the framework of the local molecular tumour board.

## Material & Methods

### Histology and immunohistochemistry analyses

Tissue was fixed with 4 % formaldehyde and embedded in paraffin. Using a microtome, 5 μm thin sections were cut, deparaffinized and rehydrated in water. Hematoxylin and eosin (HE) staining was carried out on a Tissue Tek Prisma Plus autostainer (SAKURA).

For immunohistochemistry, sections were stained with antibodies directed against Paired Box Gene 8 (PAX8), GATA Binding Protein 3 (GATA3), Trefoil Factor 1 (TFF1), Erb-B2 Receptor Tyrosine Kinase 2 (ERBB2/HER2), Oestrogen-receptor (ESR1/ER), Progesterone-receptor (PGR/PR), WT1 Transcription Factor (WT1), Tumour Associated Calcium Signal Transducer 2 (TACSTD2/TROP2), pan-neurotrophic receptor tyrosine kinase (NTRK), Programmed Death Ligand 1 (PD-L1/CD274), Calbindin 2/Calretinin (CALB2), Inhibin alpha (INHA), Thyroglobulin (TG), Synaptophysin (SYN), Membrane Metalloendopeptidase (MME/CD10), PMS1 Homolog 2, Mismatch Repair System Component (PMS2), MutL protein homolog 1 (MLH1), MutS Homolog 6 (MSH6), and MutS Homolog 2 (MSH2). All antibodies used in this study have been validated and controlled for specificity using internal and external positive and negative controls. For the TROP2 analysis, a histoscore (H-score) with a range from 1-300 was calculated based on the percentage of stained cells and the staining intensity (H-score = 1 x % weakly positive cells + 2 x % moderately positive cells + 3 x % strongly positive cells). Antigen retrieval was achieved with ER1 (EDTA-buffer Bond pH 6.0; 20 minutes) or ER2 (EDTA-buffer Bond pH 9.0; 20 minutes). IHC was carried out on the autostainer BOND RX system (Leica Biosystems). The immunoreaction was visualized with the Bond Polymer Refine Detection Kit (DS 9800; brown labelling; Novocastra; Leica Biosystems) resulting in a brown colour and counterstained with hematoxylin. The stained tissue sections were digitalized using a NanoZoomer S60 Digital slide scanner (Hamamatsu) at 40 x magnification. The antibodies are summarized in **Supplementary table 3**.

### Organoid establishment and maintenance

Primary tumour cells were obtained from advanced-stage ovarian cancer patients undergoing routine surgery (Ethical Approvals 563/21 & 631/24). Patient data is summarized in **Supplementary table 4**. To isolate primary cells, tumour resections were incubated with 1 U/μl collagenase I (Sigma) for 2 hours. Ovarian organoids were established following protocols from Trillsch, Czogalla (7). Digested cells were washed with DPBS (PanBioTech) spun down at 300 x g at 4°C for 5 min, and directly seeded in Matrigel. After 30 min, Matrigel domes were overlaid with ovarian cancer organoid medium (Advanced DMEM/F-12 (Gibco) supplemented with 10 mM HEPES (Capricorn), 1% GlutaMax (Gibco), 1% Pen/Strep (Biochrom), 1x NCS21 (Capricorn), 1x N2 (Capricorn), 10 ng/mL human EGF (Peprotech), 1.25 mM N-acetyl-L-cysteine (Sigma), 0.5 μM TGF-β receptor kinase Inhibitor IV (A83-01, Cayman Chemicals), 10 μM ROCK inhibitor (Y-27632, Cayman Chemicals), 10 mM nicotinamide (Sigma), and 10 ng/ml BMP2 (QKine)). Initial passage medium was supplemented with 10 μg/ml Gentamicin (UKSH Clinics) and 1x Fungin (InvivoGen) to avoid contamination. Cells were monitored regularly during medium changes every 7 days until organoid growth was observed.

Healthy fallopian tube organoids were established from routine surgical procedures (Ethical Approval 631/24) (8, 9). Patients with pathogenic conditions of the female reproductive tract, such as malignancies or endometriosis, were not considered for organoid generation. Fallopian tube organoids were propagated in FT expansion medium (Advanced DMEM/F12 supplemented with 25% conditioned Wnt3A medium (L-Wnt3a; ATCC CRL-2647), 25% R-Spondin-1-conditioned medium, 10 mM HEPES, 1% GlutaMax, 2% B27 or NCS21, 1% N2, 10 ng/mL human recombinant EGF, 100 ng/mL human Noggin (Peprotech), 100 ng/mL human FGF10 (QKine), 1mM nicotinamide, 10 μM Y-27632, and 1 μM A83-01).

All organoids were maintained by regular passage every 7-14 days as described in Holthaus, Hedemann (8). For this, cells were harvested by scratching of Matrigel domes with a P1000 pipet tip and collected into a 0.1% BSA-coated 15 ml tube with Advanced DMEM/F-12. Cells were spun down at 500 x g at 4°C for 5 min, supernatant was removed, and cells were incubated for 5-7 min with TripLE Express (Gibco). Afterwards, organoids were mechanically disrupted by up-and-down pipetting with a syringe with 18G gauge. Then, cells were washed with Advanced DMEM/F-12, pelleted, resuspended in Matrigel and seeded into a new 24-well plate. Medium was exchanged 1-2x per week. Organoids were imaged using the NYONE Scientific (SYNENTEC). Images were extracted using YT Software (SYNENTEC) and FIJI (10).

### Chemotherapeutic treatments and viability assays

Eight wells of ovarian cancer organoids were harvested from 24-well routine cultures per 96-well treatment plate. After mechanical and enzymatic disruption as described for passaging, cells were seeded in 10 μL domes into Costar Flat White Clear Bottom 96-well plates (Corning). Cells were grown for another four days and subsequently treated and imaged. For treatment, cells were incubated with binimetinib (0.1-20 μM, Cayman Chemicals), olaparib (1-100 μM, Cayman Chemicals), paclitaxel (10-500 nM), and cisplatin (0.1-20 μM, both UKSH clinics). After four days of incubation, supernatant was removed and the CellTiter-Glo 3D Cell Viability Assay (Promega) was performed. After equilibration at RT, the reagent was pipetted into the wells. Well contents were mixed and allowed to stabilize at RT for 120 min. Afterwards, luminescence was detected using a plate reader (Sparks, Tecan).

### Whole exome sequencing and bioinformatic analysis

Whole exome sequencing data was analysed using the MIRACUM pipeline (11). The pipeline detects and annotates somatic single nucleotide variants, Insertion-Deletions (InDels), and copy number variations. Furthermore, microsatellite instability (MSI), tumour mutational burden (TMB), tumour signatures, and Homologous Recombination Deficiency (HRD) scores are calculated. Mutational output was additionally used for a Circos plot that was plotted in R (12) using *circlize* (13).

### Statistical analysis

Basic calculations were performed using MS EXCEL 2019 (Microsoft). Figures were plotted using Prism 10.6.0 (GraphPad) and R Studio 4.4 (12). IC_50_ determinations were performed in Prism. The panel composition and annotations were created using Affinity Designer 2.6.0 (Serif). No power calculation was performed in this study.

## Results

### Clinical presentation and management

A 61-year-old female patient presented with a mild discomfort in the lower abdomen and increased perspiration for a duration of three weeks. Other additional symptoms were not reported. The most recent gynaecological examination was conducted seven years prior. The patient’s medical history revealed no instances of cancer in her family, no pre-existing conditions, and a history of nulligravida status.

Vaginal ultrasound and CT scan revealed a 10 × 9 × 9 cm mass originating from the left ovary (**Figure 1A**). Additionally, a capsule-transgressing lesion was identified in the spleen, raising concern for distant metastasis, and several potentially metastasized paracolic lymph nodes (**Figure 1B, C**). Furthermore, transvaginal ultrasound revealed an 11 cm inhomogeneous pelvic mass, comprising both cystic and solid components (**Figure 1D**). Further investigation using laboratory analyses revealed elevated levels of tumour markers, with cancer-antigen 19-9 (CA19-9) recorded at 168 kU/l and cancer-antigen (CA125) at 88.2 kU/l. The results of carcinoembryonic antigen and other blood tests were unremarkable.

**Figure 1:**
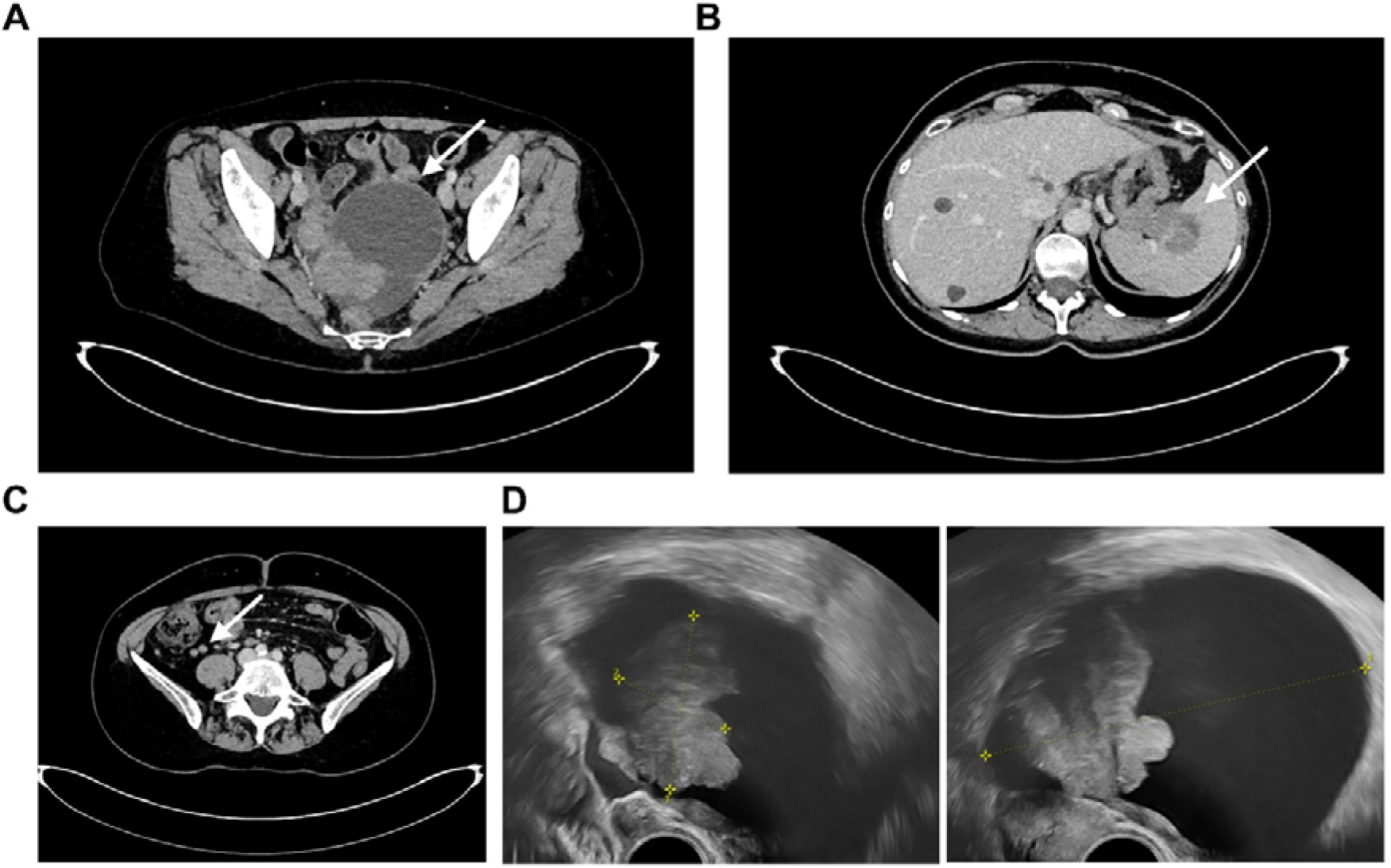
Clinical imaging suggesting an advanced ovarian malignancy with possible splenic involvement. **A** Computed tomography cross-sectional imaging of the pelvis. Large contrast-enhancing partially solid, partly cystic mass most likely originating from the left ovary, measuring approximately 10 × 9 × 8 cm. The solid portion measures 5.5 cm. **B** A predominantly hypodense mass growing beyond the spleen capsule is visible at the splenic hilum, measuring approximately 4.3 × 3.8 cm axially. **C** Two lymph nodes suspected of metastasis paracolic on the right, measuring 10 × 7 mm and 7 × 7 mm. **D** Transvaginal ultrasound report: 11 × 8 cm cystic, inhomogeneous ovarian tumour with a solid portion measuring 6 × 4 cm on the right, little free fluid in the Douglas, no ascites.

Given the findings suggestive of advanced ovarian malignancy with possible splenic involvement, the patient underwent an exploratory laparotomy. During surgery, a tumour originating in the left ovary was found to occupy the entire minor pelvis. The right adnexa were unremarkable. Only several subserosal myomas were found on the uterus, which was otherwise morphologically unremarkable. The peritoneum, the colonic fossae, and the greater omentum were palpated as tumour-free. The mesentery of the large and small intestine was also unremarkable. A total abdominal hysterectomy, bilateral salpingo-oopherectomy, omentectomy, splenectomy, and partial peritonectomy were performed. Intraoperative frozen section analysis was conducted in order to verify the clinical suspicion of ovarian cancer, which was confirmed, albeit of an uncertain histological type at that time. No macroscopical residuals (R0) was achieved. The patient recovered well from the surgery and was able to start adjuvant chemotherapy promptly. Currently there is no evidence of disease.

The postoperative histopathological examination revealed a mesonephric-like adenocarcinoma measuring 50 mm in size in the left ovary. Immunohistochemical analysis demonstrated the characteristic profile of this rare entity and are presented in **Supplementary table 1**. The tumour was found to be mismatch-repair proficient (MMRp), indicating intact DNA-repair mechanisms and therefore unlikely to respond to immune checkpoint inhibitors. A subsequent histopathological examination of the spleen revealed metastatic involvement, thereby resulting in FIGO stage IVb classification. Molecular testing including BRCA1/2 DNA Repair Associated genes (BRCA1/2) mutation analysis and homologous recombination deficiency (HRD) testing was initiated to guide targeted therapy decisions, with results pending at the time of treatment initiation.

Given the rarity of mesonephric-like ovarian adenocarcinoma and the absence of specific evidence-based treatment guideline for this histological subtype, the decision to proceed with standard first-line chemotherapy for advanced ovarian cancer was made with the patient’s agreement. The patient was started on a regimen comprising 6 cycles of carboplatin (AUC 5), paclitaxel (175 mg/m^2^) and bevacizumab (15 mg/kg) every three weeks. This was followed by maintenance therapy with bevacizumab for a total of 15 weeks. Treatment tolerance has been acceptable and the patient is currently completing her planned six cycles of chemotherapy.

### Molecular characterisation

Following the surgical removal of the patient’s tumour mass, samples were analysed by WES and IHC. IHC analysis revealed positive signals for Paired Box Gene 8 (PAX8), Trefoil Factor 1 (TFF1), Erb-B2 Receptor Tyrosine Kinase 2 (ERBB2/HER2), Tumour Associated Calcium Signal Transducer 2 (TACSTD2/TROP2), PMS1 Homolog 2/Mismatch Repair System Component (PMS2), MutL protein homolog 1 (MLH1), MutS Homolog 6 (MSH6), and MutS Homolog 2 (MSH2), focal weak to moderate positivity for GATA Binding Protein 3 (GATA3), luminal positivity for Membrane Metalloendopeptidase (MME/CD10), and questionable weak positivity for Inhibin alpha (INHA)(**Figure 2A, Supplementary Figure 1**). The tumour was negative for Oestrogen-receptor (ER/ESR1), Progesterone-receptor (PR/PGR), WT1 Transcription Factor (WT1), pan-neurotrophic receptor tyrosine kinase (NTRK), Programmed Death Ligand 1 (PD-L1/CD274), Calbindin 2/Calretinin (CALB2), Thyroglobulin (TG), and Synaptophysin (SYN) matching earlier reports of mesonephric-like adenocarcinomas (**Figure 2A, Supplementary Figure 1**) (14). Expression of DNA mismatch repair proteins (MLH1, PMS2, MSH2, MSH6) was detectable by IHC proving MMR-proficiency. 20 % of the tumour cells were negative for TROP2 while 10 % were weakly, 20 % moderately and 50 % strongly positive. More than 50 % of the tumour cells were strongly positive for ERBB2/HER2. 0 % of tumour cells showed membranous expression of PD-L1. Less than 1% of the tumour surface was covered by PD-L1-positive immune cells.

**Figure 2:**
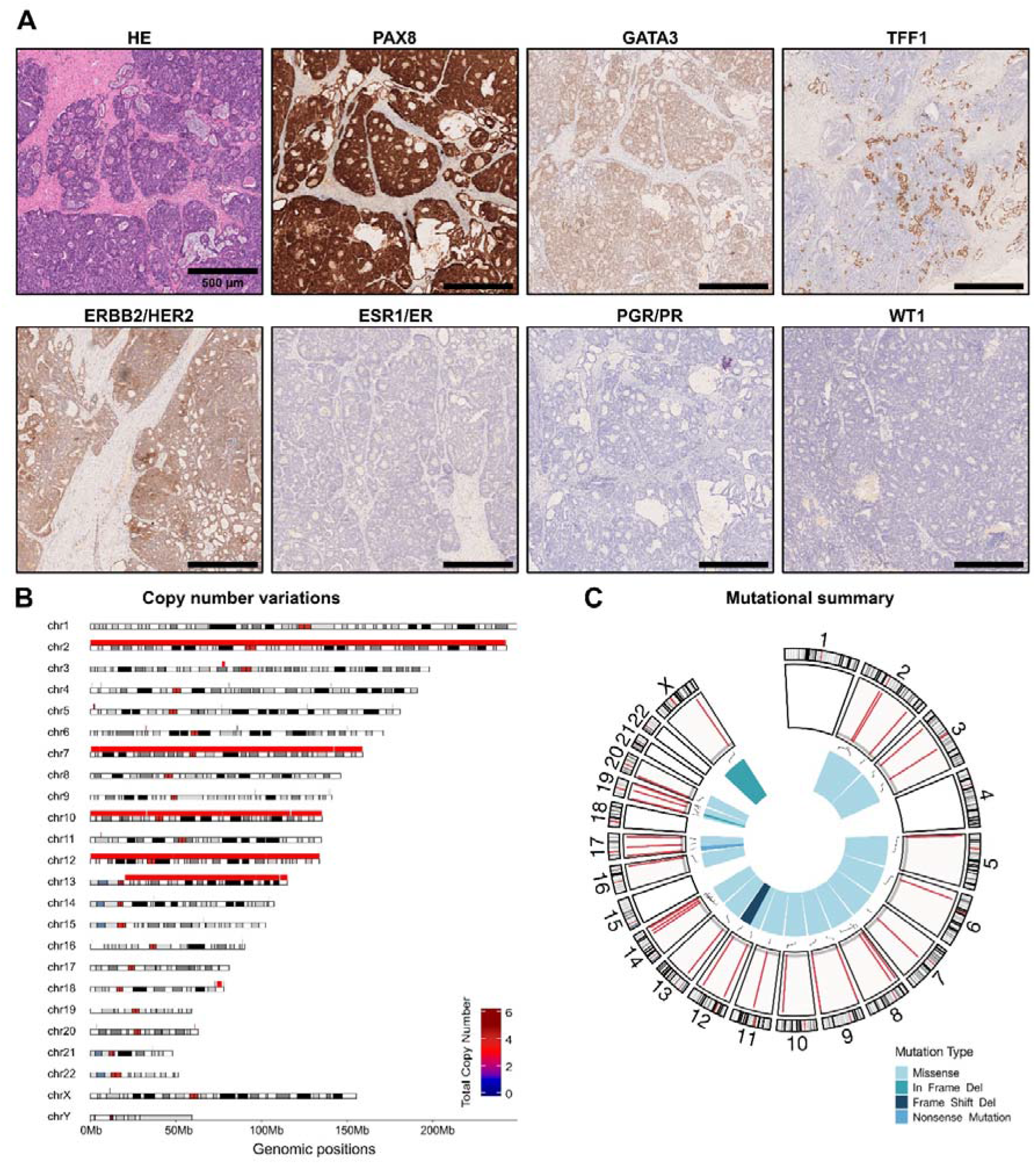
Molecular characterisation of the tumour by immunohistochemistry and whole-exome sequencing. **A** Immunohistochemical stainings of Paired Box Gene 8 (PAX8), GATA Binding Protein 3 (GATA3), Trefoil Factor 1 (TFF1), Erb-B2 Receptor Tyrosine Kinase 2 (ERBB2/HER2), Oestrogen-receptor (ER/ESR1), Progesterone-receptor (PR/PGR), and WT1 Transcription Factor (WT1). The tumour shows detectable expression of PAX8, TFF1, GATA3, ERBB2/HER2 and is negative for PR, ER, and WT1 showing characteristic marker gene expression for MLA (14). Additional stainings were performed for Tumour Associated Calcium Signal Transducer 2 (TACSTD2/TROP2), pan-neurotrophic receptor tyrosine kinase (NTRK), Programmed Death Ligand 1 (PD-L1/CD274), Calbindin 2/Calretinin (CALB2), Inhibin alpha (INHA), Thyroglobulin (TG), Synaptophysin (SYN), Membrane Metalloendopeptidase (MME/CD10), PMS1 Homolog 2/Mismatch Repair System Component (PMS2), MutL protein homolog 1 (MLH1), MutS Homolog 6 (MSH6), and MutS Homolog 2 (MSH2) and are provided in **Supplemental Figure 1. B** Overview of the copy-number variations identified by whole-exome sequencing via the MIRACUM pipeline (11). Large amplifications are found in chromosome 2, 7, 10, 12, and 13. **C** Mutational summary variations identified by whole-exome sequencing via the MIRACUM pipeline (11). A total of 31 mutations were found of which 2 were in-frame deletion, 1 was a frameshift-deletion, 1 nonsense mutation, and 27 were missense mutations.

For molecular diagnostics, whole-exome sequencing (WES) was performed on resected tumour samples. Large amplifications were found in chromosome 2, 7, 10, 12, and 13 (**Figure 2B**). Sequencing revealed 31 mutations, of which 2 were in-frame deletion, 1 was a frameshift-deletion, 1 nonsense mutation, and 27 were missense mutations (**Figure 2C, Supplementary table 2**). Two mutations, KRAS G12V and DYNC1H1 G4072S have been annotated as pathogenic and likely pathogenic by the in-house bioinformatic pipeline (11). Tumour mutational burden was low with 1.14/Mb.

### *In vitro* chemosensitivity testing

In parallel to the molecular characterisation of the tumour, ovarian cancer organoids were established from tumour samples following earlier established protocols (7, 15). The organoids showed high organoid-forming efficacies (**Figure 3A**) and stayed proliferative in culture for > 3 passages or > 4 weeks. After establishment of the *in vitro* organoid line, we conducted determinations of chemosensitivity and -resistance.

**Figure 3:**
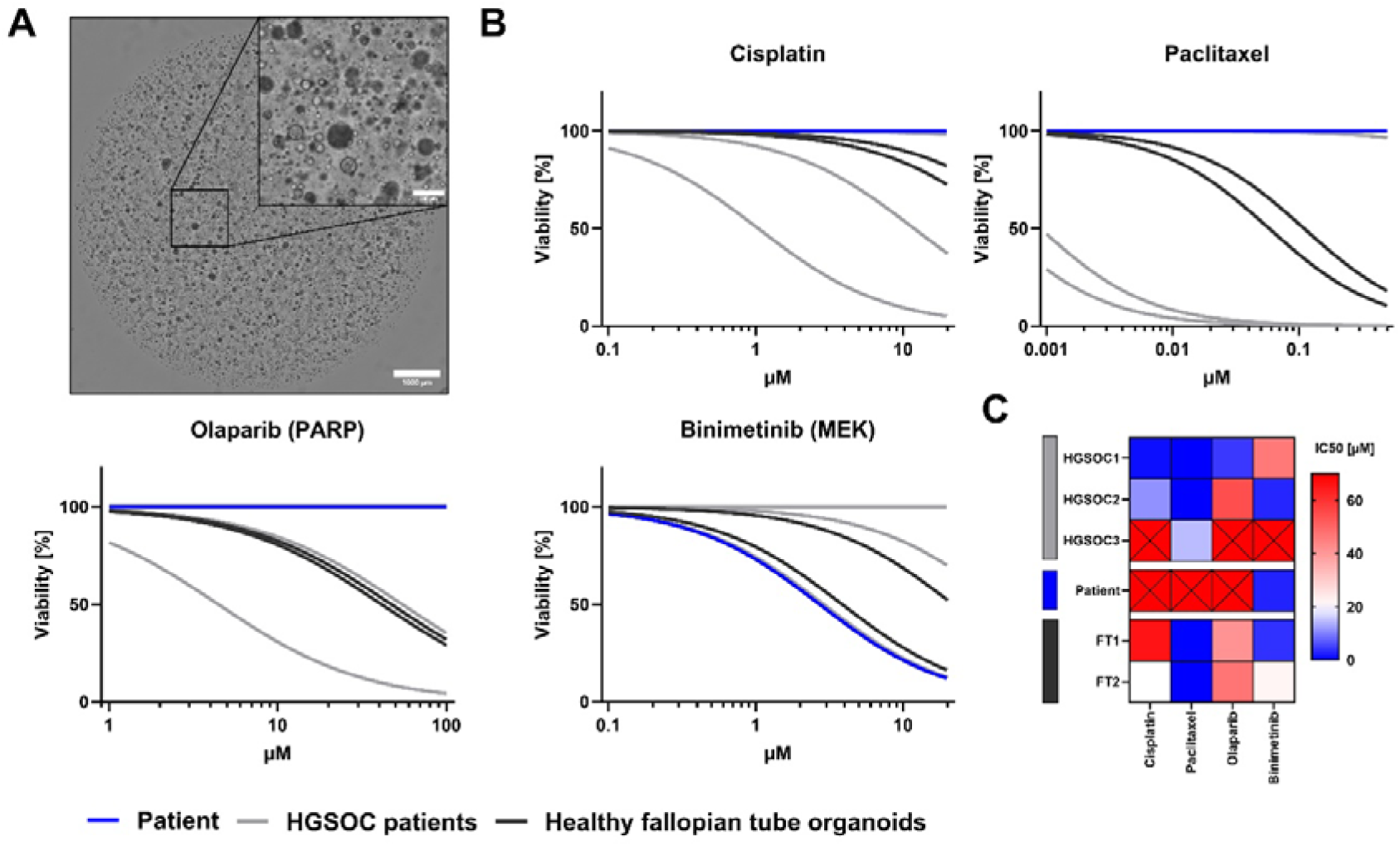
Establishment and *in vitro* chemosensitivity testing of organoids from the tumour sample. **A** Representative light microscopic images of the established ovarian cancer organoid cultures. **B** Determination of chemotherapeutic sensitivity profiles against standard therapy of platin-based chemotherapy, paclitaxel, the PARP-inhibitor olaparib, and the MEK-inhibitor binimetinib of the discussed MLA patient (Curves in blue). Three high-grade ovarian cancer (HGSOC1-3, light grey) organoid lines are displayed for comparison. Fallopian tube (FT1-2, dark gey) organoid cultures were used as toxicity controls. Parentheses indicate target proteins. **C** Comparison of IC_50_s for chemotherapeutics indicate high resistance against every tested compound except for binimetinib. Crosses indicate conditions where IC_50_ values could not be determined due to flat dose–response curves. Scale bars in **A** represent 1000 μm (overview) and 200 μm. Brightfield images were taken using a NYONE Scientific imager and extracted using YT Software 24.7 (both SYNENTEC). Curve fitting and calculation of IC_50_s were performed in GraphPad Prism 10.4.0.

For this, earlier established fallopian tube organoids were used as toxicity control (8, 9) (**Figure 3B, dark grey**) and for better comparison, three organoid lines of cases of high-grade ovarian cancer were included in the analysis (**Figure 3B, light grey**). HGSOC1 and 2 were relatively sensitive to the tested chemotherapeutics, while HGSOC3 was resistant to all them (**Figure 3B & C**). Fallopian tube organoids were used to indicate the therapeutic windows below the toxic range of chemotherapy. Cells were seeded into 96-well plates, allowed to settle and grow for another 3 days and were then treated with a titration of standard chemotherapeutics cisplatin, paclitaxel, and *Poly (ADP-Ribose) Polymerase 1* (PARP) inhibitor olaparib that is recommended for treatment in BRCA1/2 DNA Repair Associated (BRCA1/2) mutated patients. Additionally, we have included the *Mitogen-Activated Protein Kinase Kinase 1* (MEK/MAP2K1) inhibitor binimetinib as other have earlier reported on the sensitivity of mesonephric-like adenocarcinoma to MEK-inhibition in case of KRAS-mutations (16). Fallopian tube organoids showed intermediate resistance to chemotherapy to all tested compounds with IC_50_s > 20 μM for cisplatin, > 60 nM for paclitaxel, > 40 μM for olaparib, and > 4 μM for binimetinib. The MLA patient organoid line (**Figure 3B & C, blue**) showed high resistance to standard chemotherapeutics with an IC_50_ > 100 μM for cisplatin, paclitaxel, and olaparib. Only, MEK inhibition with binimetinib resulted in significant cytotoxicity as it showed a comparably low IC_50_ with 2.711 μM in comparison to other chemotherapeutics and patients.

## Discussion

Mesonephric adenocarcinomas of the ovary have only been recently classified as a new rare type of cancers of the female reproductive system (6). Due to their novelty and their rare occurrence, protocols for the generation of in vitro cellular models that would enable their use in larger studies examining targeted, personalized treatment schemes have been absent so far.

The here described tumour being PAX8^+^, GATA3^+^, TFF1^+^, ER^-^, PR^-^, WT1^-^ exhibited characteristic expression of marker proteins. Copy number variation analysis by WES showed amplifications in chromosomes 2, 7, 10, 12, and 13. Others report that MLAs are mainly driven genetically by alterations in KRAS/NRAS that also correspond to our observations (14). Especially, KRAS G12 mutations appear to be a common identifier (6, 14, 17-19). Furthermore, others have been describing similar mutational profiles in cases of mesonephric adenocarcinomas like *APC Membrane Recruitment Protein* (AMER) and BCL6 Corepressor (BCOR) mutations (20) but those appear generally non-recurring mutations. As in our case, MLAs appear to be microsatellite-stable with low tumour mutational burden (14).

Here, we describe for the first time the feasibility of organoids of MLAs for the generation of *in vitro* chemosensitivity data using established methods for the generation of HGSOC organoids (7, 9). Importantly, the *in vitro* chemosensitivity testing using the organoid model aligned well with the molecular profiling and existing literature. Downstream inhibition of KRAS signalling using the MEK-inhibitor binimetinib appeared to be highly advantageous in comparison with standard platin- and taxane-based chemotherapy. Additionally, the PARP1 inhibitor olaparib also appeared to be ineffective. These results corresponded to the associated sequencing data that showed that the tumour proved mismatch-repair proficiency and no mutations in BRCA1/2.

The here described analyses by sequencing, immunohistochemistry, and chemotherapeutic sensitivity testing revealed chemotherapeutic resistance at molecular and functional level. Despite the unfavourable profile, the patient remains disease-free to this date, although the risk of recurrence remains and requires regular clinical monitoring.

## Conclusions

The here described study is, to our knowledge, the first study utilizing organoids of mesonephric-like adenocarcinomas of the ovary for *in vitro* chemosensitivity testing. We show the applicability of existing protocols for HGSOC organoid generation. The established MLA organoids exhibit exceptional resistance to standard chemotherapy, such as cisplatin and paclitaxel, however, is sensitive to MEK inhibition. This study underlines the importance and feasibility of personalized *in vitro* models for functional precision medicine purposes also for rare entities of ovarian cancer.

## Declarations

## Abbreviations

MLA: mesonephric-like adenocarcinoma
FIGO: Fédération Internationale de Gynécologie et d’Obstétrique
SOC: standard-of-care
HGSOC: High-grade serous ovarian cancer
FT: Fallopian tube
CA19-9: cancer antigen 19-9
CA125: cancer antigen 125/ MUC16
MMRp: mismatch-repair proficient
IHC: immunohistochemistry
WES: whole exome sequencing
IC_50_: Half maximal inhibitory concentration
HE: Haematoxylin and eosin
MSI: microsatellite instability
TMB: tumour mutational burden
HRD: Homologous Recombination Deficiency

## Ethics approval and consent to participate

The studies involving humans were approved by the ethical committee of the Medical Faculty of the Christian-Albrechts-Universität zu Kiel (Ethical Approvals 563/21 & 631/24). The studies were conducted in accordance with the local legislation and institutional requirements. The participants provided their written informed consent to participate in this study.

## Consent for publication

Not applicable.

## Availability of data and materials

The datasets used and/or analysed during the current study are available from the corresponding author on reasonable request.

## Competing interests

The authors declare that they have no competing interests.

## Funding

The authors acknowledge funding from the University Hospital Schleswig-Holstein. DH, CR & NH acknowledge funding by the University Cancer Center Schleswig-Holstein via the Twinning Grant and PHSH Shared-Campus-Awards Initiative. DH & NH thank UKSH-Gemeinsam Gutes tun! Förderstiftung des UKSH (017_2023) for funds enabling ovarian organoid cultures. HDL and CR acknowledge funding via the UKSH Clinician Scientist program. FK was funded by the German Research Foundation ‘Clinician Scientist Program in Evolutionary Medicine’ (project number 413490537). The funders played no role in study design, data collection, analysis and interpretation of data, or the writing of this manuscript.

## Authors’ contributions

DH conducted organoid experiments. HDL summarized clinical outcomes. HDL and IA provided tissues. FK, TM, and BK performed pathological evaluations. DH, HDL, and LM analysed and visualized the results. CR, VW, NM, and IA supported clinical data acquisition. LB and SF supported comparison with analyses of the molecular tumour board. JPW and NM provided infrastructure and access to the UKSH gynaecological biobank via the P2N network. HB provided and analysed sequencing data. NH and DH conceptualized and supervised the project. NH initiated close collaboration with the MTB, supported by DB & MvM. DH, HDL, and CR wrote the draft manuscript, supported by NH. All authors contributed to the article and approved the submitted version.

DH and HDL contributed equally to this manuscript, and each has the right to list themselves first in author order on their CVs.

## Acknowledgements

We thank Finja Grundt, Frauke Grohmann, and Catharina Verkooyen for their excellent technical support. We thank all other involved colleagues from the Department of Gynaecology and Obstetrics at the University Hospital Schleswig-Holstein in Kiel for the provision of tissues. We thank Dr. Reinhild Geisen and Dr. Anna Willms from SYNENTEC GmbH for technical assistance with the NyOne Scientific. We acknowledge financial support by Land Schleswig-Holstein within the funding programme Open Access Publikationsfonds.

## Supplementary materials

**Supplementary figure 1:**
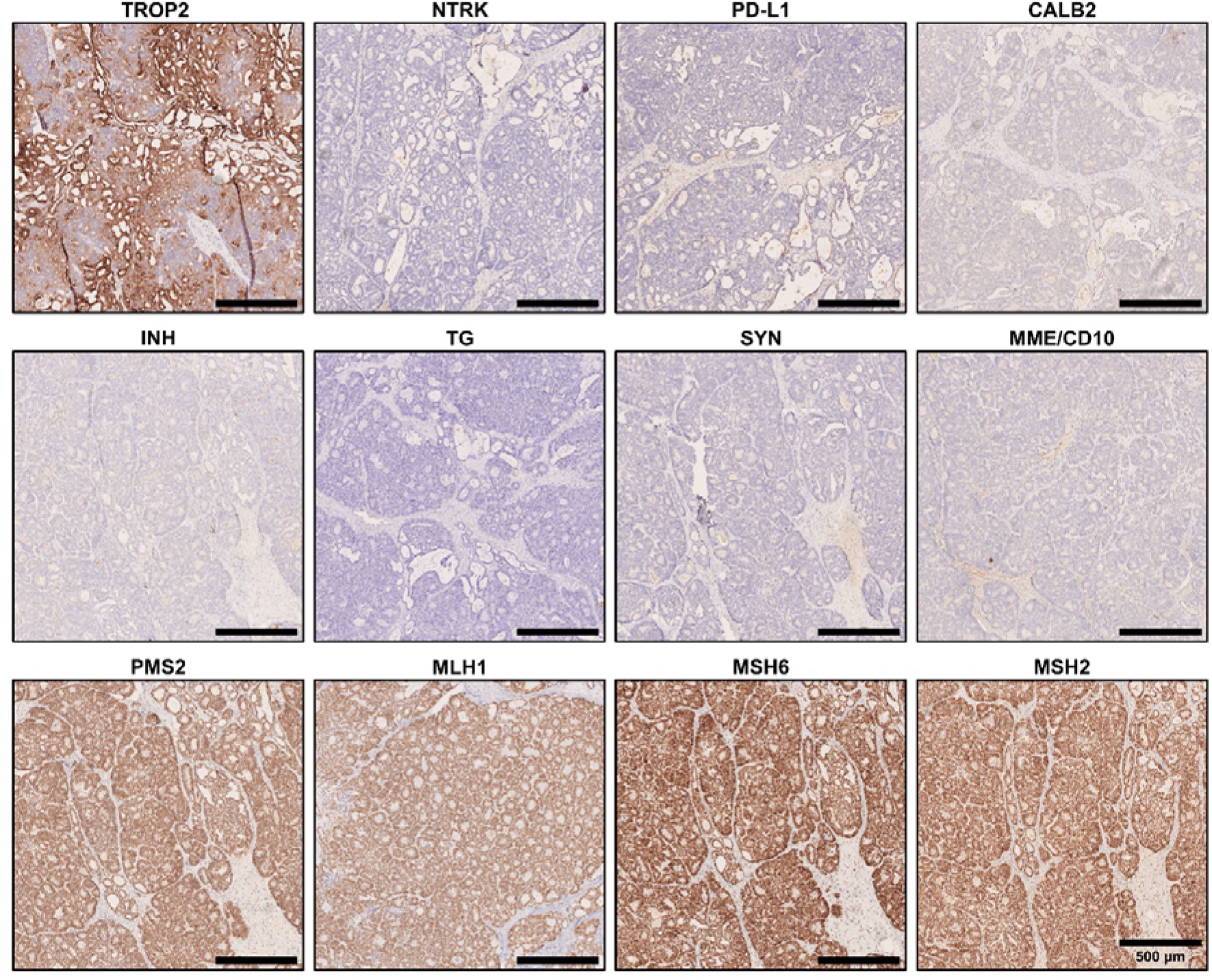
Supplementary immunohistochemistry analyses of the tumour. The tissue is positive for Tumour Associated Calcium Signal Transducer 2 (TACSTD2/TROP2), PMS1 Homolog 2/Mismatch Repair System Component (PMS2), MutL protein homolog 1 (MLH1), MutS Homolog 6 (MSH6), and MutS Homolog 2 (MSH2). Membrane Metalloendopeptidase (MME/CD10) is focally luminally positive, Inhibin alpha (INHA) focally questionable weakly positive, while pan-Neurotrophic receptor tyrosine kinase (NTRK), Programmed Death Ligand 1 (PD-L1/CD274), Calbindin 2/Calretinin (CALB2), Thyroglobulin (TG), and Synaptophysin (SYN) are negative.

**Supplementary table 1:**
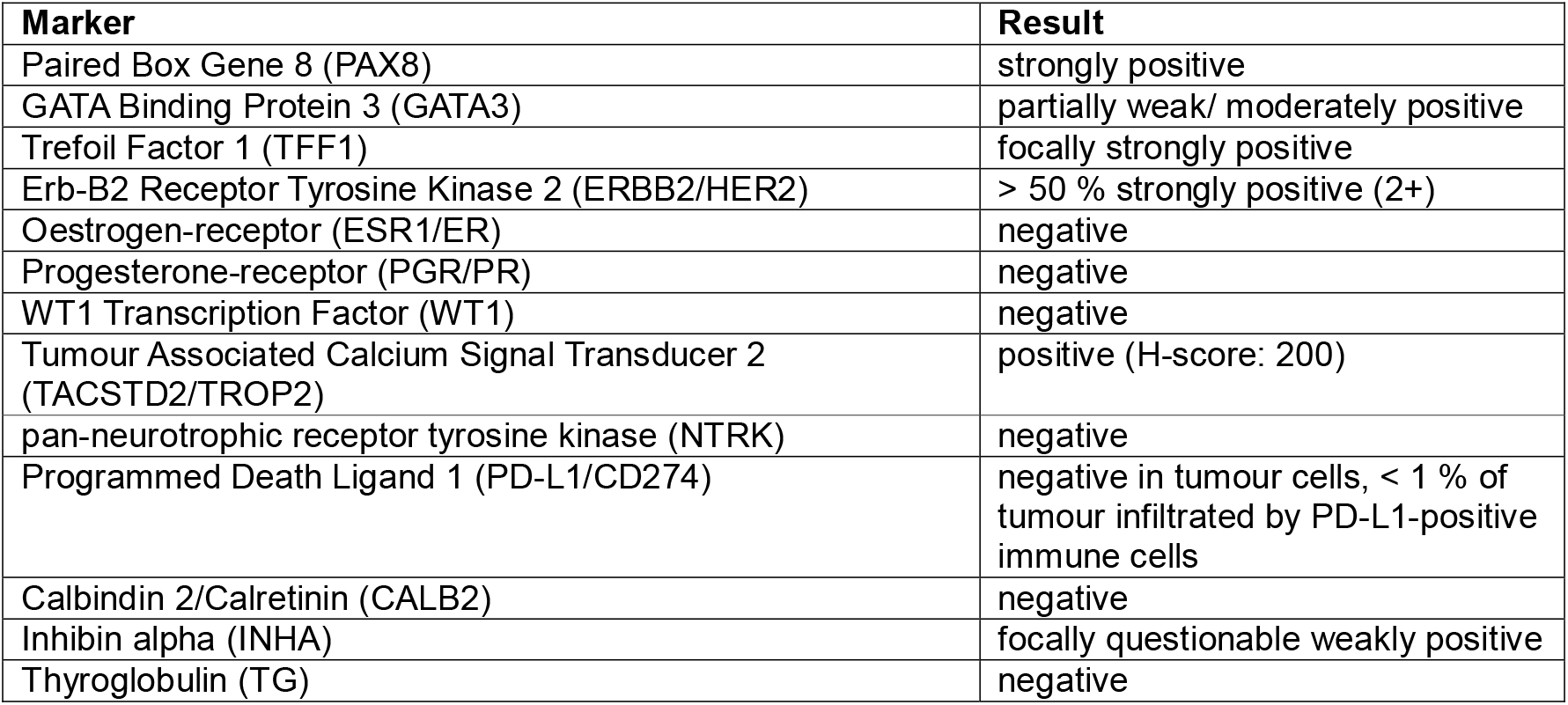
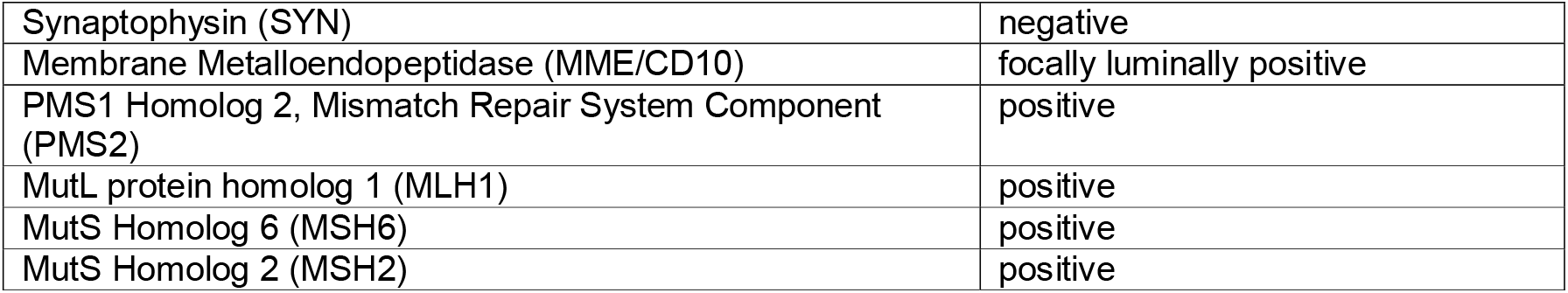
Summary of immunohistochemical analyses.

**Supplementary table 2:**
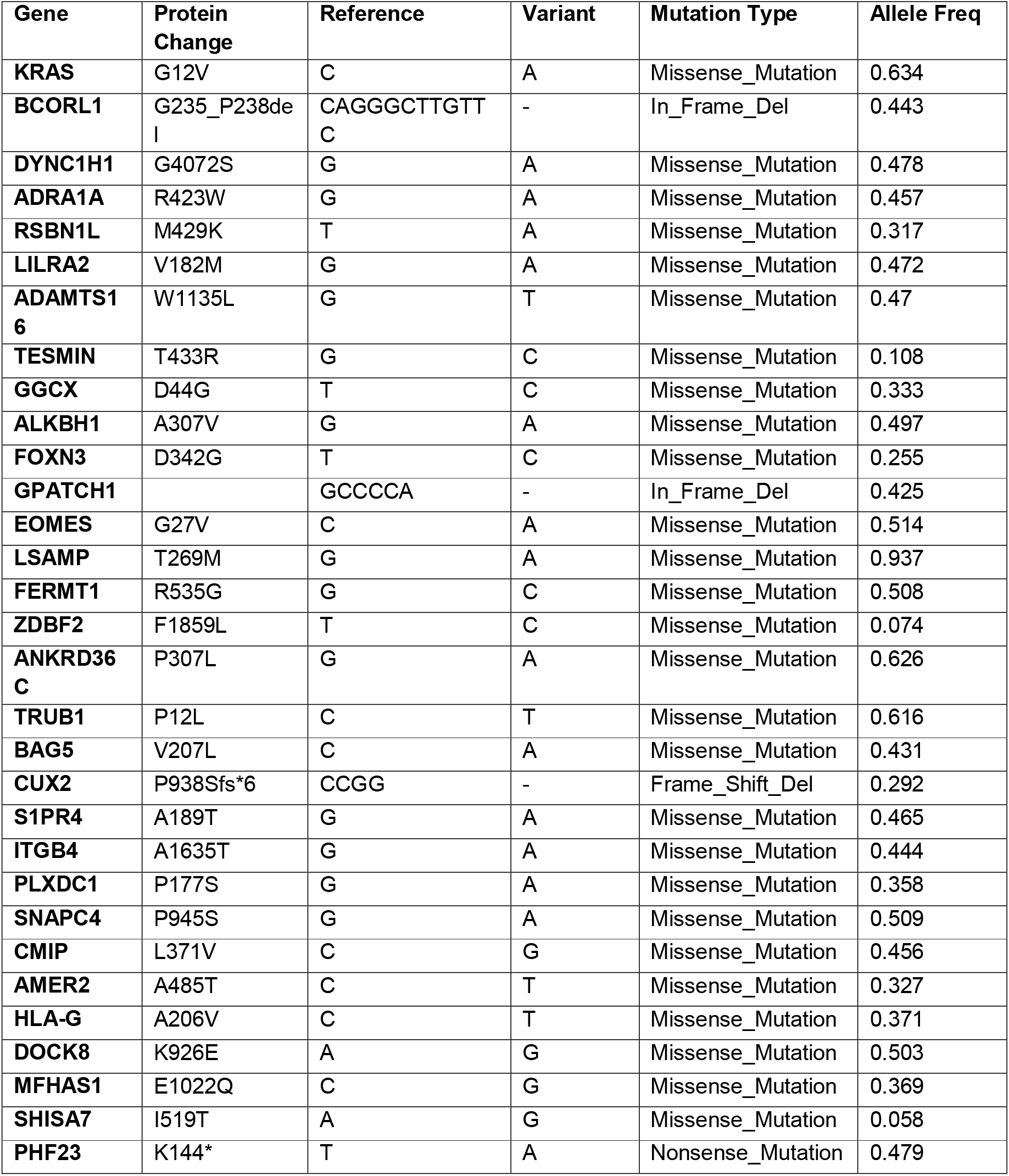
Mutational profile of the discussed patient.

**Supplementary table 3:**
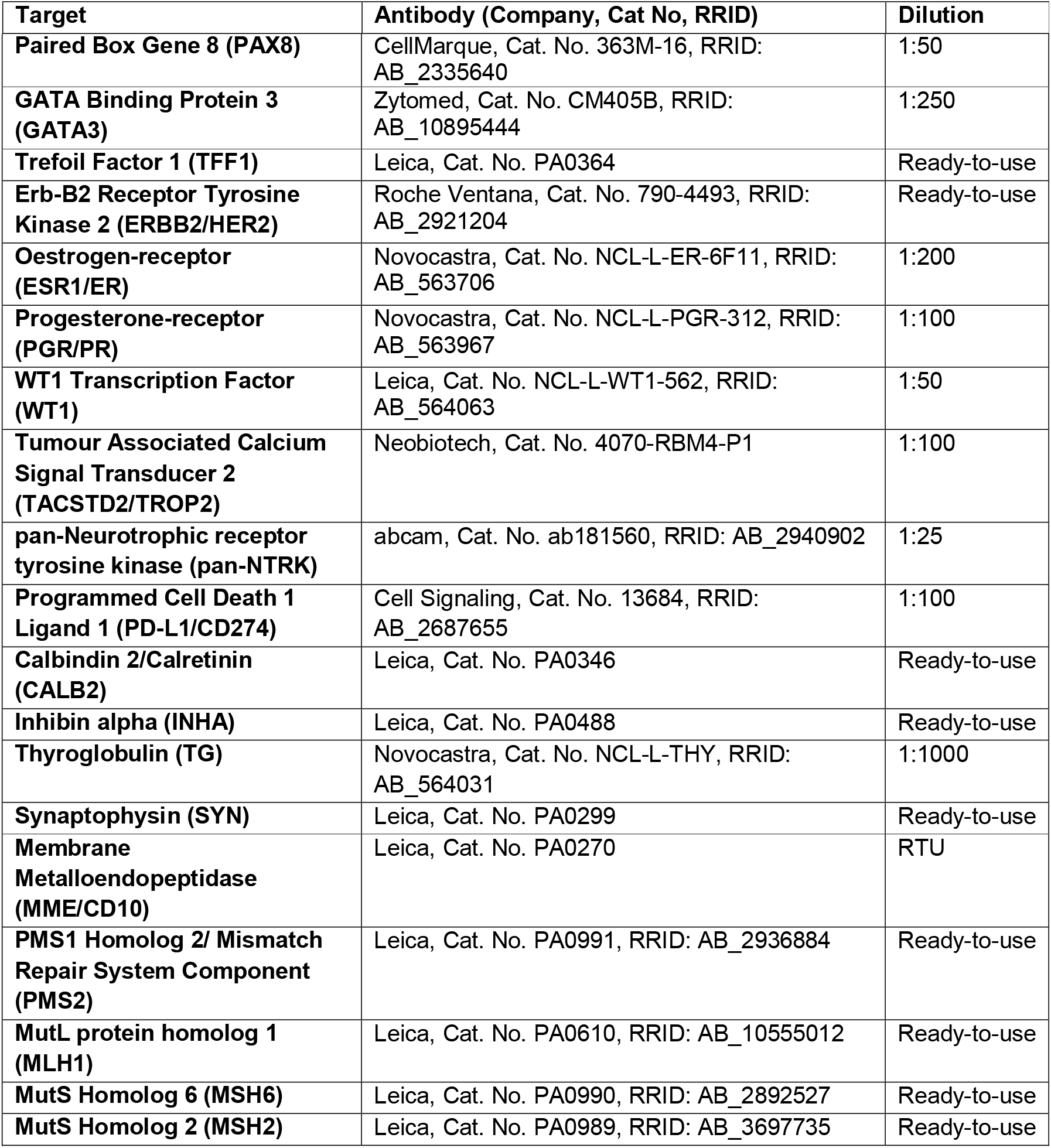
Summary of the antibodies used for IHC.

**Supplementary table 4:**
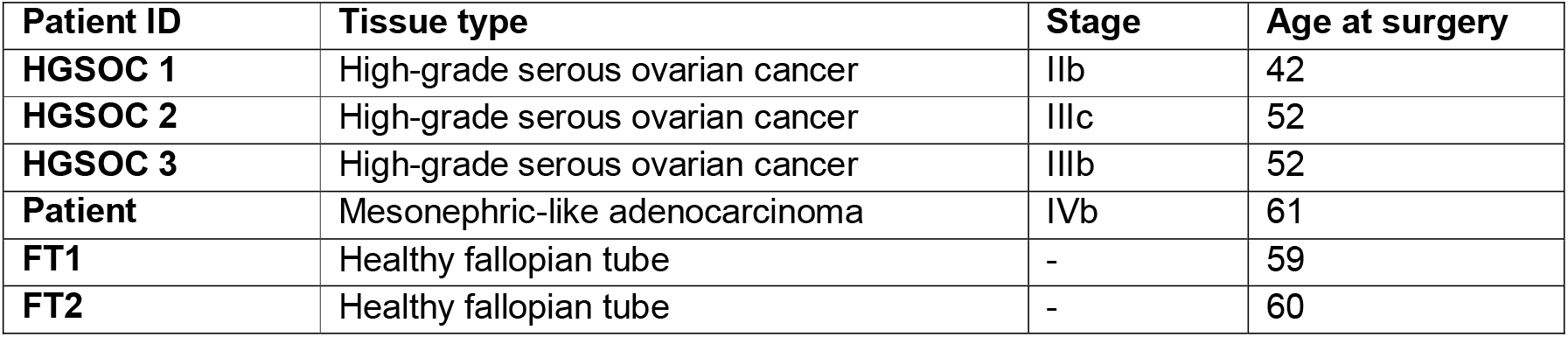
Patient characteristics.

## Notes

### Competing Interest Statement

The authors have declared no competing interest.

