## supplementary materials for "Organoids serve as viable *in vitro* model for functional precision medicine for mesonephric-like adenocarcinoma of the ovary"

* shared authorship

^#^ Corresponding author

Dr. Nina Hedemann

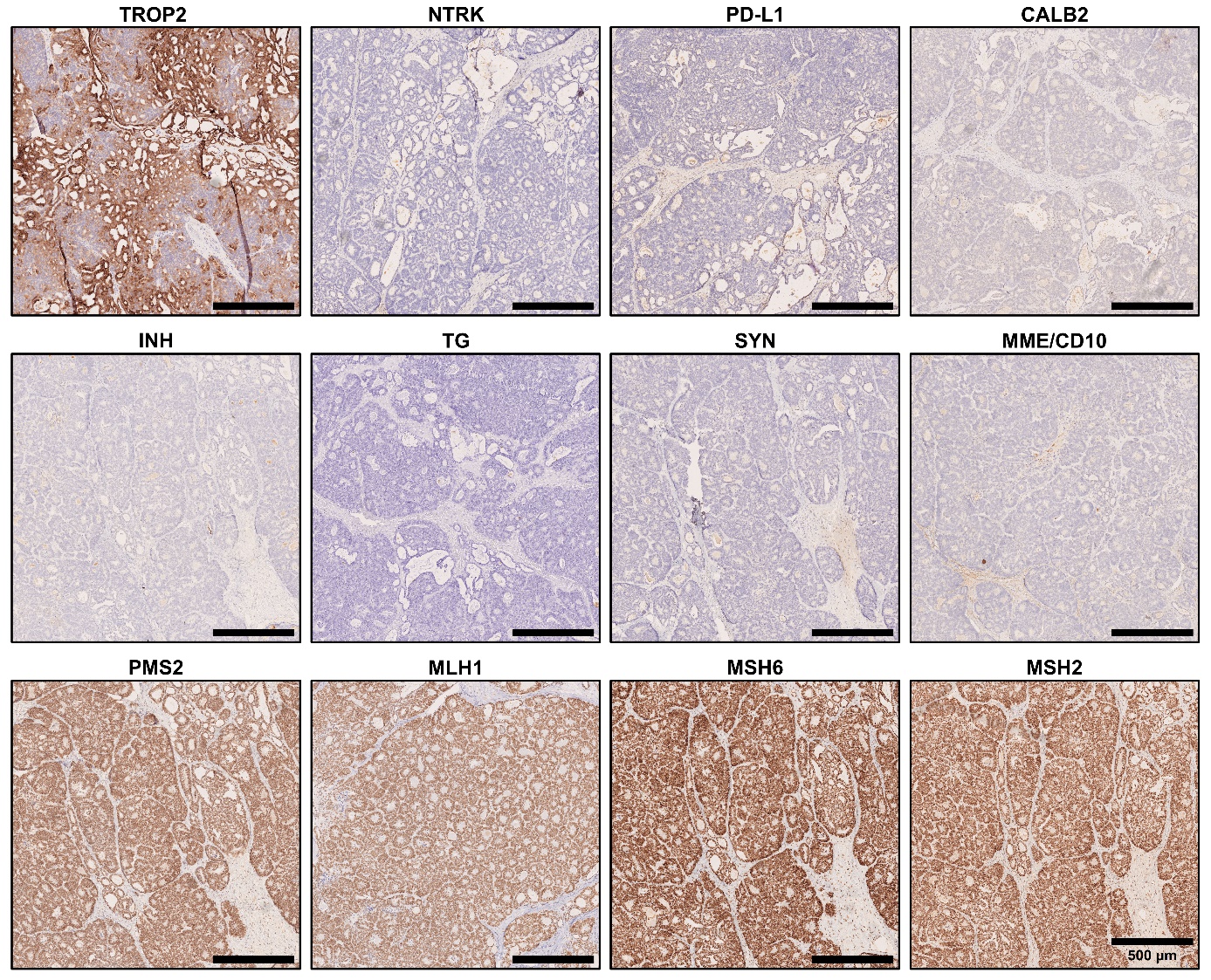

**Supplementary figure 1: Supplementary immunohistochemistry analyses of the tumour.** The tissue is positive for Tumour Associated Calcium Signal Transducer 2 (TACSTD2/TROP2), PMS1 Homolog 2/Mismatch Repair System Component (PMS2), MutL protein homolog 1 (MLH1), MutS Homolog 6 (MSH6), and MutS Homolog 2 (MSH2). Membrane Metalloendopeptidase (MME/CD10) is focally luminally positive, Inhibin alpha (INHA) focally questionable weakly positive, while pan-Neurotrophic receptor tyrosine kinase (NTRK), Programmed Death Ligand 1 (PD-L1/CD274), Calbindin 2/Calretinin (CALB2), Thyroglobulin (TG), and Synaptophysin (SYN) are negative.

**Supplementary table 1: Summary of immunohistochemical analyses.**

| **Marker** | **Result** |
| --- | --- |
| Paired Box Gene 8 (PAX8) | strongly positive |
| GATA Binding Protein 3 (GATA3) | partially weak/ moderately positive |
| Trefoil Factor 1 (TFF1) | focally strongly positive |
| Erb-B2 Receptor Tyrosine Kinase 2 (ERBB2/HER2) | > 50 % strongly positive (2+) |
| Oestrogen-receptor (ESR1/ER) | negative |
| Progesterone-receptor (PGR/PR) | negative |
| WT1 Transcription Factor (WT1) | negative |
| Tumour Associated Calcium Signal Transducer 2 (TACSTD2/TROP2) | positive (H-score: 200) |
| pan-neurotrophic receptor tyrosine kinase (NTRK) | negative |
| Programmed Death Ligand 1 (PD-L1/CD274) | negative in tumour cells, < 1 % of tumour infiltrated by PD-L1-positive immune cells |
| Calbindin 2/Calretinin (CALB2) | negative |
| Inhibin alpha (INHA) | focally questionable weakly positive |
| Thyroglobulin (TG) | negative |
| Synaptophysin (SYN) | negative |
| Membrane Metalloendopeptidase (MME/CD10) | focally luminally positive |
| PMS1 Homolog 2, Mismatch Repair System Component (PMS2) | positive |
| MutL protein homolog 1 (MLH1) | positive |
| MutS Homolog 6 (MSH6) | positive |
| MutS Homolog 2 (MSH2) | positive |

**Supplementary table 2: Mutational profile of the discussed patient.**

| **Gene** | **Protein Change** | **Reference** | **Variant** | **Mutation Type** | **Allele Freq** |
| --- | --- | --- | --- | --- | --- |
| **KRAS** | G12V | C | A | Missense_Mutation | 0.634 |
| **BCORL1** | G235_P238del | CAGGGCTTGTTC | - | In_Frame_Del | 0.443 |
| **DYNC1H1** | G4072S | G | A | Missense_Mutation | 0.478 |
| **ADRA1A** | R423W | G | A | Missense_Mutation | 0.457 |
| **RSBN1L** | M429K | T | A | Missense_Mutation | 0.317 |
| **LILRA2** | V182M | G | A | Missense_Mutation | 0.472 |
| **ADAMTS16** | W1135L | G | T | Missense_Mutation | 0.47 |
| **TESMIN** | T433R | G | C | Missense_Mutation | 0.108 |
| **GGCX** | D44G | T | C | Missense_Mutation | 0.333 |
| **ALKBH1** | A307V | G | A | Missense_Mutation | 0.497 |
| **FOXN3** | D342G | T | C | Missense_Mutation | 0.255 |
| **GPATCH1** |  | GCCCCA | - | In_Frame_Del | 0.425 |
| **EOMES** | G27V | C | A | Missense_Mutation | 0.514 |
| **LSAMP** | T269M | G | A | Missense_Mutation | 0.937 |
| **FERMT1** | R535G | G | C | Missense_Mutation | 0.508 |
| **ZDBF2** | F1859L | T | C | Missense_Mutation | 0.074 |
| **ANKRD36C** | P307L | G | A | Missense_Mutation | 0.626 |
| **TRUB1** | P12L | C | T | Missense_Mutation | 0.616 |
| **BAG5** | V207L | C | A | Missense_Mutation | 0.431 |
| **CUX2** | P938Sfs*6 | CCGG | - | Frame_Shift_Del | 0.292 |
| **S1PR4** | A189T | G | A | Missense_Mutation | 0.465 |
| **ITGB4** | A1635T | G | A | Missense_Mutation | 0.444 |
| **PLXDC1** | P177S | G | A | Missense_Mutation | 0.358 |
| **SNAPC4** | P945S | G | A | Missense_Mutation | 0.509 |
| **CMIP** | L371V | C | G | Missense_Mutation | 0.456 |
| **AMER2** | A485T | C | T | Missense_Mutation | 0.327 |
| **HLA-G** | A206V | C | T | Missense_Mutation | 0.371 |
| **DOCK8** | K926E | A | G | Missense_Mutation | 0.503 |
| **MFHAS1** | E1022Q | C | G | Missense_Mutation | 0.369 |
| **SHISA7** | I519T | A | G | Missense_Mutation | 0.058 |
| **PHF23** | K144* | T | A | Nonsense_Mutation | 0.479 |

**Supplementary table 3: Summary of the antibodies used for IHC.**

| **Target** | **Antibody (Company, Cat No, RRID)** | **Dilution** |
| --- | --- | --- |
| **Paired Box Gene 8 (PAX8)** | CellMarque, Cat. No. 363M-16, RRID: AB_2335640 | 1:50 |
| **GATA Binding Protein 3 (GATA3)** | Zytomed, Cat. No. CM405B, RRID: AB_10895444 | 1:250 |
| **Trefoil Factor 1 (TFF1)** | Leica, Cat. No. PA0364 | Ready-to-use |
| **Erb-B2 Receptor Tyrosine Kinase 2 (ERBB2/HER2)** | Roche Ventana, Cat. No. 790-4493, RRID: AB_2921204 | Ready-to-use |
| **Oestrogen-receptor (ESR1/ER)** | Novocastra, Cat. No. NCL-L-ER-6F11, RRID: AB_563706 | 1:200 |
| **Progesterone-receptor (PGR/PR)** | Novocastra, Cat. No. NCL-L-PGR-312, RRID: AB_563967 | 1:100 |
| **WT1 Transcription Factor (WT1)** | Leica, Cat. No. NCL-L-WT1-562, RRID: AB_564063 | 1:50 |
| **Tumour Associated Calcium Signal Transducer 2 (TACSTD2/TROP2)** | Neobiotech, Cat. No. 4070-RBM4-P1 | 1:100 |
| **pan-Neurotrophic receptor tyrosine kinase (pan-NTRK)** | abcam, Cat. No. ab181560, RRID: AB_2940902 | 1:25 |
| **Programmed Cell Death 1 Ligand 1 (PD-L1/CD274)** | Cell Signaling, Cat. No. 13684, RRID: AB_2687655 | 1:100 |
| **Calbindin 2/Calretinin (CALB2)** | Leica, Cat. No. PA0346 | Ready-to-use |
| **Inhibin alpha (INHA)** | Leica, Cat. No. PA0488 | Ready-to-use |
| **Thyroglobulin (TG)** | Novocastra, Cat. No. NCL-L-THY, RRID: AB_564031 | 1:1000 |
| **Synaptophysin (SYN)** | Leica, Cat. No. PA0299 | Ready-to-use |
| **Membrane Metalloendopeptidase (MME/CD10)** | Leica, Cat. No. PA0270 | RTU |
| **PMS1 Homolog 2/ Mismatch Repair System Component (PMS2)** | Leica, Cat. No. PA0991, RRID: AB_2936884 | Ready-to-use |
| **MutL protein homolog 1 (MLH1)** | Leica, Cat. No. PA0610, RRID: AB_10555012 | Ready-to-use |
| **MutS Homolog 6 (MSH6)** | Leica, Cat. No. PA0990, RRID: AB_2892527 | Ready-to-use |
| **MutS Homolog 2 (MSH2)** | Leica, Cat. No. PA0989, RRID: AB_3697735 | Ready-to-use |

**Supplementary table 4: Patient characteristics**

| **Patient ID** | **Tissue type** | **Stage** | **Age at surgery** |
| --- | --- | --- | --- |
| **HGSOC 1** | High-grade serous ovarian cancer | IIb | 42 |
| **HGSOC 2** | High-grade serous ovarian cancer | IIIc | 52 |
| **HGSOC 3** | High-grade serous ovarian cancer | IIIb | 52 |
| **Patient** | Mesonephric-like adenocarcinoma | IVb | 61 |
| **FT1** | Healthy fallopian tube | - | 59 |
| **FT2** | Healthy fallopian tube | - | 60 |
